# Zooming into rearranged genome: applying pipeline of cytological, genomic, and transcriptomic methods for structural variant interpretation

**DOI:** 10.64898/2025.12.29.696952

**Authors:** Maria Gridina, Timofey Lagunov, Polina Belokopytova, Nikita Torgunakov, Artem Nurislamov, Darya A Yurchenko, Zhanna G Markova, Tatiana V Markova, Yana Stepanchuk, Galina Koksharova, Pavel Orlov, Anna Subbotovskaia, Oxana Ryzhkova, Nadezhda V Shilova, Veniamin Fishman

## Abstract

Recent advances in genomic technologies have greatly enhanced our understanding of genotype–phenotype relationships and improved diagnostic of genetic diseases. However, the dissection of complex structural variants remains challenging due to the limitations of current methods in resolving their breakpoint and interpreting phenotypes involving multiple disrupted genes. In this study, we demonstrate how an integrative approach—combining cytological, genomic, and transcriptomic methods—enables the detection, structural and functional characterization of a complex structural variants affecting the *MBD5*, *USP34*, and *XPO1* genes. Our findings underscore the utility of the Exo-C, a modified chromosome conformation capture technique, in resolving complex rearrangements. We also report, for the first time, a composite neurodevelopmental phenotype resulting from the combined effects of *MBD5*-associated intellectual disability and 2p15p16.1 microdeletion syndromes.

## Introduction

Genetic variation spans a wide spectrum, ranging from single-nucleotide variants (SNVs) and small insertions or deletions (INDELs) to larger structural variants (SVs), defined as genomic alterations exceeding 50 base pairs. SVs include deletions, duplications, inversions, insertions, and translocations, and collectively contribute significantly to human disease^1^. Multiple genomic technologies have been developed to detect SVs, each with distinct advantages and limitations in sensitivity, resolution, and variant type detection^2^. Among the most widely used methods are conventional karyotyping, array comparative genomic hybridization (aCGH), short-read whole-exome (WES), and whole-genome sequencing (WGS). Karyotyping reliably detects large, megabase-scale SVs but lacks the resolution to precisely characterize breakpoints. aCGH is a cost-effective approach for identifying copy number changes but cannot define breakpoint sequences or detect balanced rearrangements such as inversions and translocations. WES is limited to the exome and poorly detects SVs with breakpoints outside coding regions. While short-read WGS offers genome-wide coverage and the potential to detect a broad spectrum of SVs, its cost precludes routine use in clinics, and its sensitivity is reduced in repetitive regions.

In addition to the aforementioned, other methods can provide clinically relevant insights into structural variants. Transcriptome analysis, for example, enables the detection of chimeric transcripts, which can indicate translocation events and gene fusions^3^. Chromatin conformation capture techniques, such as Hi-C, were originally developed to study 3D genome organization, highlighting its role in gene regulation^4^ and disease^5,6^ but have since been repurposed for structural variant detection and chromosomes assembly^7, 8^. Building on the efficiency of Hi-C in identifying SVs, we recently developed Exo-C^2^ – a genomic protocol that integrates chromosome conformation capture with exome enrichment. This method enables comprehensive detection of clinically significant variants, including balanced rearrangements often missed by conventional exome sequencing. Exo-C provides a cost-effective alternative to standard WES, offering improved resolution of chromosomal rearrangements and serving as a powerful tool for resolving karyotypic abnormalities of uncertain significance.

Congenital neurodevelopmental disorders (NDD) are complex conditions that can be caused by multiple genomic alternations. Due to the genetic heterogeneity, molecular diagnosis of neurodevelopmental phenotypes is challenging, and low diagnostic rate in the cohorts of NDD patients^9^ indicates that both detection and functional interpretation of the variants needs to be improved.

In this study, we demonstrate, using a representative and diagnostically challenging case, how an integrative pipeline combining cytogenetic, genomic, and transcriptomic methods can be applied to resolve and functionally interpret a complex structural variant in a patient with neurodevelopmental disorder of previously unknown etiology. We show that while karyotyping can indicate the presence of a complex structural variant, it lacks resolution to identify casual submicroscopic events near rearrangement breakpoints. Exo-C provides higher sensitivity for submicroscopic variant detection and allows the development of an inexpensive targeted Oxford Nanopore sequencing (ONT) assay to fully resolve breakpoints. Provided with the structure of chromosomal rearrangements, it is possible to utilize iPS cell models to study its transcriptional consequences. Altogether, these results enable to resolve association between complex structural variant and previously undescribed combination of clinical features caused by simultaneous disruption of *MBD5, USP34,* and *XPO1* genes.

## Materials and methods

### Human samples

Patient samples were collected from authorized medical centers. The study was approved by the local ethics committee of the Institute of Cytology and Genetics (protocol number 17, 16.12.2022). An Informed Consent was obtained from the parents included in the study. Blood samples were collected as described earlier^10^.

### Cytogenetic and Molecular Cytogenetic Studies

Cytogenetic analysis of the patient and her parents were performed on GTG-banded metaphase spreads obtained from cultured peripheral blood lymphocytes according to standard procedures. FISH was carried on cultured peripheral blood lymphocytes with DNA probes for centromeric regions of chromosome 2 and chromosome 4 (SE 2 (D2Z1); SE 4 (D4Z1)), for the short and long arms of chromosomes 2 and 4 (Arm Specific Short Probe 2, Arm Specific long Probe 2; Arm Specific Short Probe 4, Arm Specific long Probe 4) (KREATECH, The Netherlands) following the manufacturers’ protocols. FISH-analysis was performed using AxioImager M.1 epifluorescence microscope (Carl Zeiss, Jena, Germany) and an Isis digital image processing software (MetaSystems, Altlussheim, Germany).

### Chromosomal microarray

The CytoScan HD array (Affymetrix, Santa Clara, USA) was applied to detect the CNV across the entire genome following the manufacturer’s protocols. Microarray-based copy number analysis was performed using the Chromosome Analysis Suite software version 4.0 (Thermo Fisher Scientific Inc.) and are presented according to the International System for Human Cytogenomic Nomenclature 2020 (ISCN, 2020).

Detected CNVs totally assessed by comparing them with published literature and the public databases: Database of Genomic Variants (DGV) (http://dgv.tcag.ca/dgv/app/home, accessed on 31 January 2022), DECIPHER (http://decipher.sanger.ac.uk/, accessed on 31 January 2022) and OMIM (http://www.ncbi.nlm.nih.gov/omim, accessed on 31 January 2022). Genomic positions refer to the Human Genome February 2009 assembly (GRCh37/hg19), accessed on 31 January 2022. The pathogenicity of variants evaluated according to the American College of Medical Genetics (ACMG) standard guidelines^11^.

### Homemade DNA Probe

Homemade DNA probes for chromosome 2 (2p15) were used to characterize the structure of derivative chromosomes 2. The homemade DNA probe consisted of two overlapping oligonucleotide fragments of about 10 kb each.

Primers for the unique sequences of gene XPO1 (2p15 region) was selected using the standard bioinformatics tools (BLAST), which are provided by the National Center for Biotechnological Information of the United States (NCBI)^12^ and UCSC genome browser database (http://genome.ucsc.edu). Locus-specific DNA products were synthesized using long-range PCR by the BioMaster LR HS-PCR (2x) (BIOLABMIX LLC, Novosibirsk, Russia) on a GeneAmp PCR System 9700 (Applied Biosystems, California, USA) according to the manufacturer’s protocols (http://www.biolabmix.ru). All data are presented in Supplementary Table 1. The derived oligonucleotides were labeled with Green 496 dUTP (the unique sequences of gene XPO1) (Enzo Life Sciences, Inc., NY, USA) by nick-translation and denatured at 75°C for 7 min. Chromosome slides were preincubated in 2SSC at 37°C for 45 min, denatured in 70% formamide/2SSC at 73°C for 3 min, and then dehydrated at −20°C in ethanol. Probe-hybridization mixture was applied on the chromosomes, incubated at 37°C for 16 h. Slides were washed three times in 4SSC, 0.1% Tween 20 at 45°C. The data were analyzed using the epifluorescence microscope AxioImager M.1 (Carl Zeiss, Jena, Germany) and the Isis software (MetaSystems, Altlussheim, Germany).

### 3C experiment

The 3C part was performed on the peripheral blood mononuclear cells (PBMC) according to DNase I protocol^10^. In brief, 2.5 million cells were fixed using 1% formaldehyde, lysed, and then subjected to digestion with DNase I (Thermo Fisher Scientific, USA). The digested DNA ends were labeled with biotin-15-dCTP (Biosan, Russia) and ligated overnight using T4 DNA ligase (Thermo Fisher Scientific, USA). To reverse the formaldehyde crosslinks, the samples were incubated with Proteinase K (NEB, UK) at 65°C overnight, followed by ethanol precipitation. Unligated biotin-labeled ends were removed through treatment with T4 polymerase (NEB, UK). The KAPA HyperPlus kit (Roche) was used for NGS library preparation. Biotin-filled DNA fragments were pulled down using Dynabeads MyOne Streptavidin C1 beads (Thermo Fisher Scientific, USA), and the libraries were subsequently amplified. Exome enrichment was carried out using the KAPA HyperExome Probes and KAPA HyperCap kit (Roche), following the manufacturer’s protocol.

### Induced pluripotent stem cells generation

Reprogramming of patient blood cells into induced pluripotent stem (iPS) cells was performed by episomal vectors cocktail contained: reprogramming vectors expressed OCT4 (addgene #41813), MYC and LIN28 (addgene #41855), p53 carboxy-terminal dominant-negative fragment (addgene #41856), SOX2 and KLF4 (addgene #41814), EBNA1 (addgene # 41857), according to the previously described method^13^. Briefly, 5 × 10⁵ peripheral blood mononuclear cell were transfected with 1 μg of each vector using the Neon Transfection System (Thermo Fisher Scientific, USA) under optimized conditions (1650 V, 10 ms, 3 pulses).

The transfected cells were then plated onto mitomycin C-treated CD-1 mouse embryonic fibroblasts and cultured overnight in complete StemPro™-34 medium supplemented with cytokines (Thermo Fisher Scientific, USA). The following day, the medium was replaced with N2B27 medium (DMEM/F-12 with HEPES, 1% N-2 Supplement, 2% B-27™ Supplement, 1% GlutaMAX-I, 1% MEM NEAA, 1% Pen-Strep, and 0.1 mM 2-mercaptoethanol) containing 100 ng/mL bFGF (Thermo Fisher Scientific, USA), which was refreshed daily. On day 9, the medium was changed to iPS cell medium (DMEM/F12, 20% KnockOut Serum Replacement, 1% GlutaMAX-I, 1% MEM NEAA, 1% Pen-Strep, and 0.1 mM 2-mercaptoethanol) supplemented with 100 ng/mL bFGF. By day 16, colonies displaying typical iPS cell morphology were manually picked and expanded in iPS cell medium containing 10 ng/mL bFGF. The cells were maintained at 37°C in 5% CO₂ and passaged mechanically at a 1:5 split ratio^14^.

### ONT breakpoints sequencing

Nanopore sequencing was performed as described earlier^2^. Briefly, libraries for WGS and adaptive sampling enrichment were prepared following the standard SQK-LSK109 kit protocol (Oxford Nanopore, UK). Cas9-enriched libraries were generated using the nCATS protocol^15^, with minor modifications for Exo-C^2^. Primers used for sgRNA synthesis are listed in Supplementary Table 1. Sequencing and live basecalling were performed on a GridION device using FLO-MIN106D flow cells (Oxford Nanopore, UK). Read alignment to the hg19 reference genome was carried out with Minimap2 (v2.26)^16^.

### Transcriptome analysis

For transcriptome analysis, total RNA was extracted from the two independent clones of patient iPS cells and 5 control iPS cells obtained from healthy donors and subjected to the stranded oligo-dT-based RNA-sequencing. RNA-seq reads were aligned to the human reference genome (GRCh37/hg19) using STAR (v2.5.2b)^17^. Transcript quantification was performed with salmon (v0.14.1) in alignment-based mode. Count data was imported into R using tximport (v1.30.0) and analysed using DESeq2 package (v4.5).

### Single nucleotide variants calling and annotation

SNV calling, annotation, and filtration were performed as described previously^18^. Reads were aligned to the human reference genome (hg19) using BWA-MEM algorithm (BWA v.0.7.17)^19^. PCR-duplicates were removed with PicardTools^20^. Base recalibration and haplotype calling were performed using GATK v.3.3^21^. SNVs were annotated using ANNOVAR^22^ software and filtered based on the following criteria: genotype quality score > 20, maximum population allele frequency < 0,01%, not reported as benign in ClinVar or Leiden Open Variation Database.

After filtering, the pathogenicity of each variant was assessed according to the recommendations of the American College of Medical Genetics and Genomics (ACMG) and the Association for Molecular Pathology^23^.

### Hi-C data processing

The “fastq” files with sequences were preprocessed with cutadapt ver-4.6. Next, the juicer ver-1.5.6 was used to make the files with aligned read pairs. The “mcool” files were made by cooler ver-0.10.2 and the further analysis were made using cooltools ver-0.7.1. All analyses were performed based on the human hg19 genome build.

Compartmentalization. We computed a compartment score (CS) per 50 kb bin as 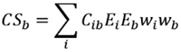, where C_ib are interchromosomal contacts between bins *i* and *b*; *E_i* and *E_b* are the first eigenvector values from the control Hi-C map at 50 kb resolution; and *w_i*, *w_b* are ICE weights after *cooler* iterative correction. The sum excludes the queried chromosome, sex chromosomes, and chromosomes harboring rearrangements (chr2 and chr4).

TAD boundaries. We estimated the insulation score using *cooltools* at 2 kb binning with a 200 kb window. Analyses were performed across SV-defined chromosomal segments ≥10 Mb (segment identifiers as in Supplementary Table 2).

### Automatic CNV detection

Exo-C data were aligned to the hg19 human genome build using BWA (v 0.7.17)^19^. BAM file sorting and indexing was performed using samtools (v 1.6)^24^. We used the following tools for CNV detection based on sequence alignments: CNVkit (v 0.9.10)^25^, CoNIFER (v 0.2.2)^26^, GATK (v 4.4.0.0)^27^. All tools were used with default parameters. CNVs reported by the three tools were intersected to keep the most robust predictions. We used DGV Gold database^28^ to exclude predictions that are possibly related to population CNV data and are not sample-specific.

### Allele-Specific Expression Analysis

Exo-C sequencing data were aligned to the human reference genome (hg19) using BWA v0.7.17. BAM files were sorted and indexed with samtools v1.6, and PCR duplicates were marked using Picard v2.27.5 (MarkDuplicates)^20^. RNA-seq reads were aligned to hg19 with STAR v2.5.2b^17^.

SNVs were identified from both RNA-seq and Exo-C alignments using GATK v4.6.1.0, retaining all positions including those with zero alternative counts in RNA-seq. Multiallelic sites were split using bcftools norm (v1.22). Exo-C heterozygous SNVs were filtered to retain sites with minimum sequencing depth of 20, genotype quality ≥20, and variant quality ≥30. Corresponding RNA-seq SNVs were also filtered for minimum depth and genotype quality.

Concordance between RNA-seq replicates was assessed using two criteria: (1) the difference in B-allele frequency (BAF) between replicates was ≤0.10, and (2) the difference in deviation of RNA-seq BAF from Exo-C BAF between replicates was ≤0.10. Additionally, Exo-C allele counts were required to be balanced (allele frequency between 0.30 and 0.70).

Allele-specific expression at each SNV was assessed by comparing RNA-seq allele counts to Exo-C counts using two-sided Fisher’s exact tests and binomial tests. Each RNA-seq replicate was tested separately against Exo-C counts, and for the combined test, allele counts from both replicates were summed before statistical testing. P-values were adjusted for multiple testing using Benjamini-Hochberg FDR correction to obtain q-values.

All analyses were performed in Python (numpy v2.3.5^29^, pandas v2.3.3^30^, scipy v1.16.3^31^). Genome-wide results were visualized with Manhattan-style plots using matplotlib v3.10.0^32^.

## Results and Discussion

### Case report

The male proband was born at 40 weeks of gestation from an eighth pregnancy (the fourth, sixth, seventh pregnancies were early miscarriages, the other four children are healthy) in a non consanguineous family. The delivery was normal, with an APGAR score of 8/9, a birth weight of was 3,300 g, and a length of 52 cm. The proband reached early motor and speech milestones with a delay — he could sit with kyphotic spine, has been walking since 1 year 2 months but had broad-based, unsteady gait, had no words. At the age of 2 years, he was diagnosed with global developmental delay. Diagnostic workup also revealed a patent foramen ovale and a bicuspid aortic valve. Brain MRI revealed hypoplasia of corpus callosum, mild frontotemporal lobe atrophy and a cyst of the septum pellucidum. The patient was referred to medical geneticist according to development delay at the age of 2 years. According to physical examination the patient had broad forehead, prominent ears, a long nose, macrostomia, downturned corners of the mouth, downslated palpebral fissures, short philtrum, short upper labial frenulum, long fingers and toes with narrowed distal phalanges. The patient had light skin and hair color with atopic patches. According to neurological examination the boy didn’t pay attention to requests, didn’t speak (made only sounds), had defuse muscular hypotonia, but normal reflexes of upper and lower limbs.

### Resolving complex chromosomal rearrangement using combination of cytological and genomic methods

To study genomic variants underlyinging patient’s phenotype, we first performed aCGH and karyotyping. There were no pathogenic and likely pathogenic CNVs by chromosomal microarray analysis: arr(X,Y)×1,(1-22)×2 (Supplementary Fig. 1). G-banded karyotyping revealed a *de novo* complex translocation involving the short arms of chromosomes 2 and 4, as well as the long arms of chromosomes 2 and 4. Subsequent FISH analysis using a PCR-specific probe identified an additional inversion or insertion within the translocated material (Fig. 1a, b). To resolve details of this SV, we profiled it using Exo-C technique. Based on the Exo-C data, we did not detect any pathogenic SNVs in patient’s exome, as well as any pathogenic CNVs (Supplementary Table 3). Exo-C identified 11 chromosomal breakpoints, confirming the arrangement of chromosomes 2 and 4 identified by karyotyping, including insertion of small (∼60 Kb) fragment on the chromosome 2 (Fig. 1c, d, e). Although we note that, this insertion occurs close to the chromosomes 2-4 junction (245 kb from the g-m junction (Fig. 1g)), it is unclear whether intrachromosomal and interchromosomal rearrangements occur independently or in frame of one complex event. To ensure that small insertion is not caused by artifact of Exo-C analysis we further validated it by sequencing Hi-C library without exome enrichment (Supplementary Fig. 2a) and performing FISH with the probe specific to the inserted fragment (Fig. 1f). Both assays confirmed intrachromosomal insertion, and Hi-C additionally validate breakpoints identified by Exo-C.

**Fig. 1.**
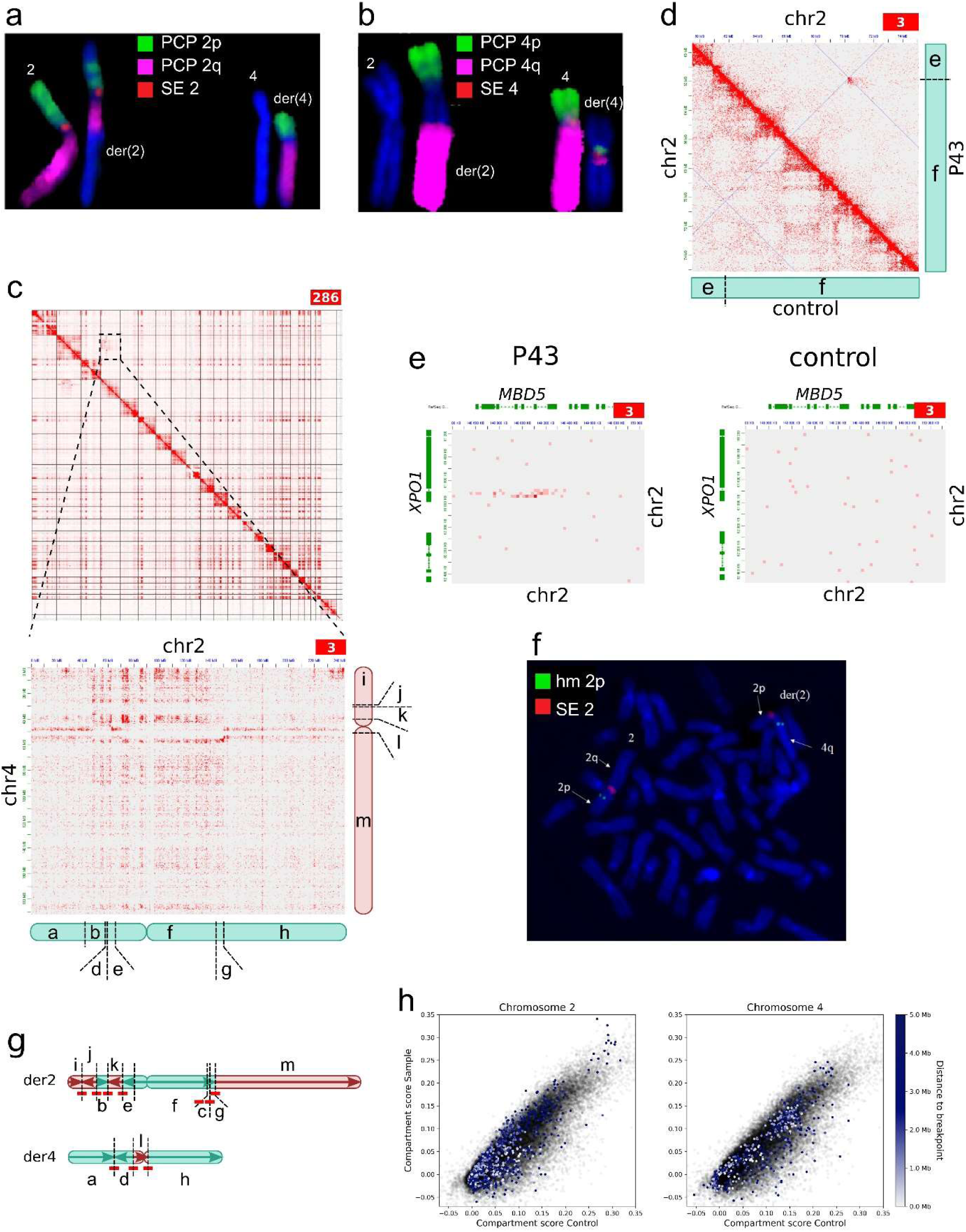
Resolving breakpoints of complex chromosomal rearrangement in patient P43. (a) FISH results performed using partial chromosome paint (PCP) DNA probes for the short arm of chromosome 2 (PCP 2p-Green), the long arm of chromosome 2 (PCP 2q-TexasRed), and the centromeric region of chromosome 2 (SE 2-Red). (b) FISH results performed with PCP DNA probes for the short arm of chromosome 4 (PCP 4p-Green), the long arm of chromosome 4 (PCP 4q-TexasRed), and the centromeric region of chromosome 4 (SE 4-Red). (c) Interchromosomal Exo-C contact maps for P43, supporting the SV structure. The letters denote the fragments resulting from chromosomal breakage. (d) Hi-C contact map of the chromosome 2 region for P43 (above the diagonal) and a control sample (with a normal karyotype; below the diagonal). The contact map indicates a breakpoint junction pattern between regions chr2:61,750,001–61,760,000 and chr2:70,220,001–70,230,000 in the patient sample. This was the only junction not confirmed by nanopore sequencing. (e) Hi-C contact map of the chromosome 2 region for P43, indicating the pattern for a small intrachromosomal insertion. The same region is shown for the control sample (with a normal karyotype). (f) FISH analysis using centromeric region of chromosome 2 (SE 2 - Red) and a custom-labeled 2p15 probe (Green) revealed insertion of the XPO1 locus (2p15) into the q23 region of the derivative chromosome 2. (g) Schematic representation of SVs detected in P43 cells based on Exo-C analysis. Arrows indicate the direction of increasing genomic coordinates. Red lines indicate breakpoints identified with single-nucleotide resolution using nanopore sequencing. (h) Scatter plots comparing compartment scores between the patient and control (50-kb bins). Semi-transparent gray points denote bins from chromosomes without rearrangements. Colored points mark bins on the rearranged chromosome that lie within 5 Mb of the nearest breakpoint; color encodes genomic distance to the breakpoint. Axes show the compartment score in the control (x-axis) versus the patient (y-axis).

Exo-C and Hi-C data described above do not detect SV breakpoints with nucleotide resolution. However, resolution of these methods was sufficient to design enrichment system for Oxford Nanopore Sequencing^15^, targeting the regions flanking breakpoints. This allows us to identify breakpoints with nucleotide resolution at low cost – with ∼0.6X average genome coverage we obtain 1-8X coverage for most of the breakpoints. However, enrichment efficiency varies from locus to locus (Supplementary Table 4), and for one out of 11 breakpoints we did not obtain sufficient coverage (Fig. 1g, Supplementary Table 2). Specifically, we were unable to confirm the breakpoint connecting XPO1 exon 4 and the intergenic region on chromosome 2. We note this is a limitation of the ONT enrichment system: although efficient in most cases, it might require optimization for a subset of loci. Thus, the combination of cytogenetic analysis and Exo-C allowed us to characterize the structure of the complex chromosomal rearrangement as:

der(2)(4pter→4p14::2p21→2p15::4p12→4p14::2p15→2p15::2p13.3→2q23.1:
:4q12→4qter),der(4)(2pter→2p21::2p15→2p13.3::4p12→4q12::2q23.1→2qter)dn

Complex SV have the potential to disrupt topologically associating domain boundaries or rewire chromatin compartments, thereby altering the regulatory environment of nearby genes. To assess whether the rearrangements identified in our patient lead to local perturbations of 3D genome organization, we quantitatively evaluated chromatin architecture in the vicinity of the breakpoints by computing compartmentalization scores and insulation indices from our Hi-C data. Compartment scores (CS) were calculated for the patient and the matched controls. Figure 1h shows patient–control scatter plots, where bins on unaffected chromosomes are displayed in semi-transparent gray, whereas bins located within ≤5 Mb of any breakpoint are colored according to their distance to the nearest junction. We did not observe any systematic deviation from the global patient–control correlation as a function of breakpoint proximity. In addition, we flagged bins within ≤5 Mb whose residuals exceeded 3σ from the fitted linear trend and examined their gene content (Supplementary Table 5); no genes with established disease associations were identified. We next compared insulation profiles between the patient and the control and computed the 3σ threshold based on the standard deviation of patient–control differences. None of the breakpoint-adjacent regions exhibited insulation changes exceeding this significance threshold (Supplementary Fig. 2b). These analyses indicate that the complex rearrangements in this case do not cause detectable compartment-level or insulation-level perturbations in the surrounding genomic regions.

To complement the 3D genome analyses and assess whether the rearrangement produces broader cis-regulatory consequences, we generated induced pluripotent stem cells from the patient’s samples and performed transcriptome analysis on two cell clones altogether with control iPS cells lines obtained from the healthy donors. We performed a differential gene expression analysis comparing the patient-derived iPS cell line to control iPS cell lines. The analysis identified 46 differentially expressed genes (FDR < 0.05) (Supplementary Fig. 3a). Nearly all of these were located on chromosomes unaffected by the complex SV. Only two genes - CXCL5 and ANKRD36 - reside on chromosome 2, at 19 Mb and 27 Mb from the nearest breakpoint, respectively (Supplementary Table 6), distances far exceeding typical ranges of SV-driven cis effects. Moreover, the genes are not OMIM-listed, and their altered expression is unlikely to contribute to the patient’s clinical presentation.

### Functional analysis of SV consequences provides explanation of NDD phenotype

We note that intrachromosomal insertion of the 60 288 bp on chromosome 2 affects coding sequences of *USP34, XPO1*, and *MBD5* genes (Fig. 1f, g, 2a). Disruption of *MBD5* gene is known to cause autosomal dominant intellectual developmental disorder-1 (MRD1) (OMIM #156200)^33^. Additionally, genes USP34 and XPO1 located in 2p15 cytoband and might contribute to the development of chromosome 2p16.1-p15 deletion syndrome^34,35,36^. The fragment inserted into the 4th exon of *MBD5* contains the terminal part of *XPO1* (including last exon and polyadenylation signal), followed by the promoter and first exon of *USP34* gene. As a result of the insertion, coding strands of all three genes are co-directional with *MBD5* and *USP34* promoters, potentially allowing their transcription. Thus, functional interpretation of the discovered structural variant depends on whether the resulting chimeric allele can encode a valid template for MBD5, *XPO1*, or *MBD5* proteins or their fragments. Confirming the structure of the rearrangement described above, transcriptome analysis revealed split reads connecting the 4th exon of *XPO1* to the genomic region chr2:70,218,973-70,220,156 (Fig. 1g, Fig. 2f, Supplementary Fig.3c), which was not observed in the ONT sequencing data. We also observed evidence of chimeric transcript including genes ASPRV1 (chr2:70,215,909-70,216,025) and MSH2 (chr2:47,652,793-47,652,942), supporting another breakpoint revealed by karyotyping, Exo-C, and ONT data.

**Fig. 2.**
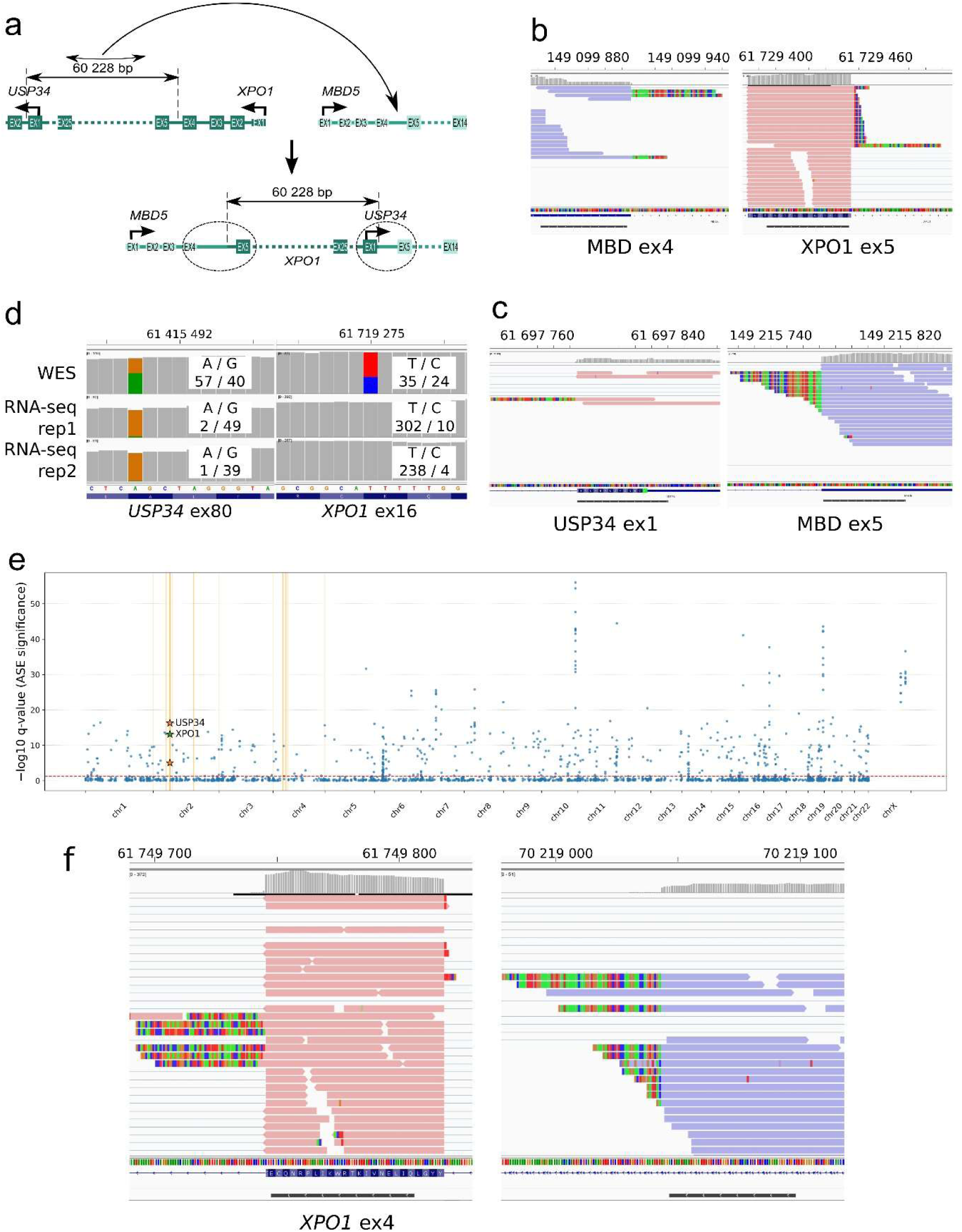
Transcriptomic validation of chromosome 2 insertion. (a) Schematic representation of the chromosome 2 insertion detected using Exo-C and Hi-C data. The fragment chr2:61,688,256–61,748,484 is inverted and inserted into chr2:149,116,886–149,116,865. The genomic breakpoints of the insertion disrupt the USP34, XPO1, and MBD5 genes. (b, c) IGV screenshots showing split reads confirming the presence of chimeric transcripts: (b) MBD5-XPO1 and (c) USP34-MBD5. Gray lines indicate BLAT alignment of soft-clipped reads. (d) IGV screenshot showing allelic representation of single-nucleotide variants in the USP34 and XPO1 exons based on Exo-C sequencing and transcriptome sequencing data. (e) Genome-wide allele-specific expression results are visualized using Manhattan-style plots generated with Matplotlib v3.10.0. The x-axis represents chromosomes, and the y-axis shows −log10(q-value), indicating ASE significance. Yellow vertical lines denote structural variant breakpoints, and stars mark genes located near the breakpoints. (f) IGV screenshot showing split reads confirming the breakpoint junction between regions chr2:61,750,001–61,760,000 and chr2:70,220,001–70,230,000. Gray lines indicate BLAT alignment of soft-clipped reads.

We next focused on transcriptomic reads aligning near 60 Kb insertion in the *MBD5* gene. This analysis identified two types of abnormal splicing products, both concordant with the proposed insertion structure. The first transcript originates from the 1st exon of *USP34* and extends into the 5th exon of *MBD5* (Fig. 2a, c). Exon 1 of *USP34* and exon 5 of *MBD5* are in different open reading frames, therefore translation initiated at the start codon of *USP34* prematurely terminates within *MBD5* exon 5 (Supplementary Fig. 3b). Thus, the fused transcript does not encode a functional MBD5 sequence and is likely targeted for degradation via nonsense-mediated RNA decay^37^.

The second transcript consists of exons 1-4 of *MBD5*, followed by a transition to exon 5 of *XPO1* (Fig. 2b). The start codons located in exon 5 of MBD5 and exon 2 of XPO1 are not included in this transcript, resulting in the absence of a correct translation initiation site. Based on these observations, we speculate that no RNA product encoding *USP34, MBD5* or *XPO1* can be transcribed from the rearranged allele. To validate this assumption, we examine heterozygous variants within the coding sequences of *USP34, MBD5* and *XPO1*. Based on genomic Exo-C data, we identified SNVs in exons 12, 69, and 80 of the *USP34* gene and exon 16 of the XPO1 gene. However, transcriptomic reads contain only one allele, indicating that these genes are expressed in a monoallelic fashion (Fig. 2d). These findings collectively confirm disruption of these genes, leading to the expression of only one functional copy of both *USP34* and *XPO1*.

To further validate the functional consequences of the rearranged allele, we examined allele-specific expression (ASE) genome-wide, across heterozygous SNVs identified in Exo-C in comparison to RNA-seq data. Filtering was applied to retain only well-supported variants with concordant RNA-seq replicates, ensuring reliable ASE measurements.

Genome-wide ASE analysis did not reveal pronounced clustering of SNVs with strong allelic imbalance near the identified breakpoints (Fig. 2e), indicating that the structural rearrangement does not induce regional ASE outside the disrupted genes. Consistent with the prior manual transcriptomic findings, SNVs within *USP34* and *XPO1* exhibited significant allelic imbalance, confirming that only one allele is expressed. *MBD5* SNVs were either absent or failed coverage and heterozygosity filters, precluding ASE assessment for this gene.

Several extreme ASE outliers were observed elsewhere in the genome. These typically corresponded to loci nearly homozygous with extremely high allele counts in RNA-seq compared to WES, expectedly producing very low p-values. On chromosome 19, high-ASE SNVs mapped to *NLRP2*^38,39^, a gene previously reported to show consistent random monoallelic expression in iPS cell lines across clonal replicates, likely reflecting stochastic or parent-of-origin effects rather than technical artifacts.

Overall, ASE analysis confirms that *USP34* and *XPO1* are expressed from a single allele, in agreement with the transcriptomic evidence. No additional genomic regions exhibited strong localized allelic imbalance near breakpoints. These results strengthen the integrative interpretation of the complex structural variant, linking the rearrangement to loss-of-function effects at the disrupted loci.

Thus, we propose that the patient’s phenotype arises from the combined effects of haploinsufficiency of *MBD5* gene, along with *USP34* and *XPO1* genes, disturbances in which appear to significantly contribute to the distinctive craniofacial abnormalities and intellectual disability observed in 2p15p16.1 microdeletion syndrome^36,40,35^. Patient P43 presents with global developmental delay, which overlap between MBD5-associated neurodevelopmental disorder (MRD1; OMIM #156200) and 2p15-p16.1 microdeletion syndrome (OMIM #612513), exept severe speech impairment, which is more specific for MRD1^33^. The structural brain abnormalities observed in the patient, however, are characteristic of 2p15p16.1 microdeletion syndrome^41^. The patient’s dysmorphic facial features—such as a broad forehead, prominent ears, and downturned corners of the mouth – are more specific for MRD1, whereas downslanting palpebral fissures, macrostomia, and a long nose are more typical of 2p15p16.1 microdeletion syndrome. The same applies to the long fingers with tapered phalanges. Thus, combination of MRD1 and 2p15p16.1 microdeletion syndrome fully explain observed clinical features.

## Conclusion

Here we demonstrate how application of (molecular)cytogenetics, genomic, and transcriptomic methods resolve complex SV associated with NDD phenotype. We note that alternative genome-wide approaches, such as optical genome mapping (OGM), have recently emerged as powerful tools for the detection of complex and balanced structural variants. Although OGM was not applied here, we consider it a promising complementary method, particularly in diagnostic settings. In the present work, however, our goal was to illustrate the added value of integrating classical cytogenetics with Exo-C–based genomic analysis and transcriptomics for structural and functional interpretation of complex rearrangements.

While cytological and transcriptomic methods are well established as tools for molecular diagnostics, here we highlight the utility of Exo-C method that helped to identify causal variant. We revealed that observed clinical manifestations originate from the cumulative effect of loss-of-function mutations in the *MBD5*, *USP34* and *XPO1*.

## Supporting information

List of primers

Breakpoints junction confirmed by nanopore sequencing

Results of CNV prediction tools

Nanopore sequencing coverage date (FC - fold change)

Regions of the genome near SV breakpoints with outlier compartment scores and the corresponding gene content

List of differentially expressed genes identified in the patient-derived iPSCs

## Data Availability

Raw human sequencing data are classified as personal information and cannot be shared, in accordance with Russian regulations on personal data protection. Processed Exo-C data is available under accession GEO: GSM8030925. Processed Hi-C data is available under accession GEO: GSM…. Processed RNA-seq data is available under accession GEO: GSM… (IDs pending; will be available in the same GSE series as GSM8030925: GSE253950)

## Acknowledgements

Cell culture was performed at the Collective Center of ICG SB RAS “Collection of Pluripotent Human and Mammalian Cell Cultures for Biological and Biomedical Research”, project number FWNR-2025-0017.

The work was partially performed using the equipment of the Center for Collective Use “Proteomic Analysis”, supported by funding from the Ministry of Science and Higher Education of the Russian Federation (No. 075-15-2021-691).

## Author Contributions

MG performed genomic experiments with help from YS; TL, PB, NT and GK performed computational analyzis; AN performed targeted nanopore sequencing; DAY, ZGM, TVM and NVS performed FISH experiments; OR, PO and AS performed NGS; DAY, ZGM, TVM and NVS enrolled samples to cohort and interpreted the patient data; VF conceived the study and supervised the work with co-supervision from NVS; VF and MG drafted the manuscript and all authors read and approved the final manuscript.

## Ethics Declaration

Ethics approval and consent to participate. The recruitment of the cohort in this study was conducted in strict accordance with the principles of the Declaration of Helsinki and the International Conference on Harmonization Good Clinical Practice (ICH-GCP) guidelines. The study was approved by the local ethics committee of the Institute of Cytology and Genetics (protocol number 17, 16.12.2022) and the local ethics committee of the Tomsk Medical Research Center (protocol number 15, 28.02.2023). Informed Consent was obtained from all patients or their parents/representatives included in the study.

