## Supplementary material for "Zooming into rearranged genome: applying pipeline of cytological, genomic, and transcriptomic methods for structural variant interpretation": List of primers

**Supplementary Table 1**

### List of primers

| Primer | Sequence (5’ – 3’) | Description |
|---|---|---|
| 2p15-1F | CCCACATGCTGTTTTCCTCTGCTAACGA | Primers for amplification of the unique sequences of region 2p15 by long-range PCR on the short arm of chromosome 2. chr2:61705893-61715828 |
| 2p15-1R | GCTGTTGGACACCCCTTTGTAATTCAGC | |
| 2p15-2F | GCTTTGCAGGCTCTCACATTTGTTTTCA | Primers for amplification of the unique sequences of region 2p15 by long-range PCR on the short arm of chromosome 2. chr2:61715827-61724892 |
| 2p15-2R | CCCCCTGATTTCCAACCCCAGACGATAA | |
| SpCas9-scaffold-R | aaaagcaccgactcggtgcc | Reverse primer for amplification of sgRNA template from scaffold sequence in PX458 plasmid (Addgene #48138) |
| P43_A1-1 | aagcTAATACGACTCACTATAGGTATTCCTCACAATTCAAAGTGTTTTAGAGCTAGAAATAGCAAGTTAA | Forward primer for amplification of sgRNA template. Includes T7 promoter sequence for in vitro transcription. |
| P43_A1-2 | aagcTAATACGACTCACTATAGGCTGAGTGTCGTACTCACCTCGTTTTAGAGCTAGAAATAGCAAGTTAA | Forward primer for amplification of sgRNA template. Includes T7 promoter sequence for in vitro transcription. |
| P43_A2-1 | aagcTAATACGACTCACTATAGGCAGTCGAATTCTAAGCAATTGTTTTAGAGCTAGAAATAGCAAGTTAA | Forward primer for amplification of sgRNA template. Includes T7 promoter sequence for in vitro transcription. |
| P43_A2-2 | aagcTAATACGACTCACTATAGGGTATTCATACCTAACATGGTTTAGAGCTAGAAATAGCAAGTTAA | Forward primer for amplification of sgRNA template. Includes T7 promoter sequence for in vitro transcription. |
| P43_B1-1 | aagcTAATACGACTCACTATAGGTGTGACCTCTGATACTGAGCGTTTTAGAGCTAGAAATAGCAAGTTAA | Forward primer for amplification of sgRNA template. Includes T7 promoter sequence for in vitro transcription. |
| P43_B1-2 | aagcTAATACGACTCACTATAGGTTTGAACTCCTGCATGAACAGTTTTAGAGCTAGAAATAGCAAGTTAA | Forward primer for amplification of sgRNA template. Includes T7 promoter sequence for in vitro transcription. |
| P43_B2-1 | aagcTAATACGACTCACTATAGGCCTTACTTAGGGTAGTGCCAGTTTTAGAGCTAGAAATAGCAAGTTAA | Forward primer for amplification of sgRNA template. Includes T7 promoter sequence for in vitro transcription. |
| P43_B2-2 | aagcTAATACGACTCACTATAGGTGGCCAAAAATAGTGAAGTCGTTTTAGAGCTAGAAATAGCAAGTTAA | Forward primer for amplification of sgRNA template. Includes T7 promoter sequence for in vitro transcription. |
| P43_C1-1 | aagcTAATACGACTCACTATAGGCTTAGGTGTCTACTTCCCTTGTTTTAGAGCTAGAAATAGCAAGTTAA | Forward primer for amplification of sgRNA template. Includes T7 promoter sequence for in vitro transcription. |
| P43_C1-2 | aagcTAATACGACTCACTATAGGTTGTTTCTATACAATATGGCGTTTTAGAGCTAGAAATAGCAAGTTAA | Forward primer for amplification of sgRNA template. Includes T7 promoter sequence for in vitro transcription. |
| P43_C1-3 | aagcTAATACGACTCACTATAGGTGACACGCCATGTTTCCTATGTTTTAGAGCTAGAAATAGCAAGTTAA | Forward primer for amplification of sgRNA template. Includes T7 promoter sequence for in vitro transcription. |
| P43_C1-4 | aagcTAATACGACTCACTATAGGTGACCACCAAACTGGCAGCTGTTTTAGAGCTAGAAATAGCAAGTTAA | Forward primer for amplification of sgRNA template. Includes T7 promoter sequence for in vitro transcription. |
| P43_C2-1 | aagcTAATACGACTCACTATAGGTAGACCTGTCTCTACCACTAGTTTTAGAGCTAGAAATAGCAAGTTAA | Forward primer for amplification of sgRNA template. Includes T7 promoter sequence for in vitro transcription. |
| P43_C2-2 | aagcTAATACGACTCACTATAGGACACGTAGGATTCATATTGAGTTTTAGAGCTAGAAATAGCAAGTTAA | Forward primer for amplification of sgRNA template. Includes T7 promoter sequence for in vitro transcription. |
| P43_C2-3 | aagcTAATACGACTCACTATAGGAGGCGAGTCTTATGCTAACTGTTTTAGAGCTAGAAATAGCAAGTTAA | Forward primer for amplification of sgRNA template. Includes T7 promoter sequence for in vitro transcription. |
| P43_C2-4 | aagcTAATACGACTCACTATAGGAACTGAGTCTTGTACTAACCGTTTTAGAGCTAGAAATAGCAAGTTAA | Forward primer for amplification of sgRNA template. Includes T7 promoter sequence for in vitro transcription. |
| P43_D1-1 | aagcTAATACGACTCACTATAGGTACGTATCATTCTTTTTTGTTTTAGAGCTAGAAATAGCAAGTTAA | Forward primer for amplification of sgRNA template. Includes T7 promoter sequence for in vitro transcription. |
| P43_D1-2 | aagcTAATACGACTCACTATAGGTGGATTGTATAGAGCATACGTTTTAGAGCTAGAAATAGCAAGTTAA | Forward primer for amplification of sgRNA template. Includes T7 promoter sequence for in vitro transcription. |
| P43_D2-1 | aagcTAATACGACTCACTATAGGTACTGAATAAGATCGAACAGTTTTAGAGCTAGAAATAGCAAGTTAA | Forward primer for amplification of sgRNA template. Includes T7 promoter sequence for in vitro transcription. |
| P43_D2-2 | aagcTAATACGACTCACTATAGGAGTCTTCTATCAAATTCAGGGTTTTAGAGCTAGAAATAGCAAGTTAA | Forward primer for amplification of sgRNA template. Includes T7 promoter sequence for in vitro transcription. |
| P43_E1-1 | aagcTAATACGACTCACTATAGGAGTTGATCATATATGATACTGTTTTAGAGCTAGAAATAGCAAGTTAA | Forward primer for amplification of sgRNA template. Includes T7 promoter sequence for in vitro transcription. |
| P43_E1-2 | aagcTAATACGACTCACTATAGGCAGTGAATTATGGTACTCTCGTTTTAGAGCTAGAAATAGCAAGTTAA | Forward primer for amplification of sgRNA template. Includes T7 promoter sequence for in vitro transcription. |
| P43_E1-3 | aagcTAATACGACTCACTATAGGCGTTGTCCTTCTCCATACATGTTTTAGAGCTAGAAATAGCAAGTTAA | Forward primer for amplification of sgRNA template. Includes T7 promoter sequence for in vitro transcription. |
| P43_E1-4 | aagcTAATACGACTCACTATAGGTTGGGCTCTGCATCCCCGAGGTTTTAGAGCTAGAAATAGCAAGTTAA | Forward primer for amplification of sgRNA template. Includes T7 promoter sequence for in vitro transcription. |
| P43_E2-1 | aagcTAATACGACTCACTATAGGATGGGTAGGATATAAGATCGTTTTAGAGCTAGAAATAGCAAGTTAA | Forward primer for amplification of sgRNA template. Includes T7 promoter sequence for in vitro transcription. |
| P43_E2-2 | aagcTAATACGACTCACTATAGGACTAACTATTACAGATCACAGTTTTAGAGCTAGAAATAGCAAGTTAA | Forward primer for amplification of sgRNA template. Includes T7 promoter sequence for in vitro transcription. |
| P43_E2-3 | aagcTAATACGACTCACTATAGGACCTCCTATAAAGAAACGATGTTTTAGAGCTAGAAATAGCAAGTTAA | Forward primer for amplification of sgRNA template. Includes T7 promoter sequence for in vitro transcription. |
| P43_E2-4 | aagcTAATACGACTCACTATAGGAATGAATTGAGTTCCTTCTGTTTTAGAGCTAGAAATAGCAAGTTAA | Forward primer for amplification of sgRNA template. Includes T7 promoter sequence for in vitro transcription. |
| P43_F1-1 | aagcTAATACGACTCACTATAGGATAGACCCTAATTGACCCCAGTTTTAGAGCTAGAAATAGCAAGTTAA | Forward primer for amplification of sgRNA template. Includes T7 promoter sequence for in vitro transcription. |
| P43_F1-2 | aagcTAATACGACTCACTATAGGCTCATTGATAGTCCCGAGTGGTTTTAGAGCTAGAAATAGCAAGTTAA | Forward primer for amplification of sgRNA template. Includes T7 promoter sequence for in vitro transcription. |
| P43_F2-1 | aagcTAATACGACTCACTATAGGGTGGTGAACTATCATCAAGTTTTAGAGCTAGAAATAGCAAGTTAA | Forward primer for amplification of sgRNA template. Includes T7 promoter sequence for in vitro transcription. |
| P43_F2-2 | aagcTAATACGACTCACTATAGGAAGGGAGCAGGATACTTTCGTTTTAGAGCTAGAAATAGCAAGTTAA | Forward primer for amplification of sgRNA template. Includes T7 promoter sequence for in vitro transcription. |
| P43_G1-1 | aagcTAATACGACTCACTATAGGATCTACCATATTACAAGCTAGTTTTAGAGCTAGAAATAGCAAGTTAA | Forward primer for amplification of sgRNA template. Includes T7 promoter sequence for in vitro transcription. |
| P43_G1-2 | aagcTAATACGACTCACTATAGGTTCTGACGACACGATTCCGTTTTAGAGCTAGAAATAGCAAGTTAA | Forward primer for amplification of sgRNA template. Includes T7 promoter sequence for in vitro transcription. |
| P43_G2-1 | aagcTAATACGACTCACTATAGGCAATACCTAAGTTGTCATGTTTTAGAGCTAGAAATAGCAAGTTAA | Forward primer for amplification of sgRNA template. Includes T7 promoter sequence for in vitro transcription. |
| P43_G2-2 | aagcTAATACGACTCACTATAGGTATAAGTCGTACCTTCCTATGTTTTAGAGCTAGAAATAGCAAGTTAA | Forward primer for amplification of sgRNA template. Includes T7 promoter sequence for in vitro transcription. |
