## Supplementary material for "Zooming into rearranged genome: applying pipeline of cytological, genomic, and transcriptomic methods for structural variant interpretation": Breakpoints junction confirmed by nanopore sequencing

**Supplementary Table 2**

### Breakpoints junction confirmed by nanopore sequencing

| Chromosome | Start | End | Chromosome | Start | End | Name | Type | Genes |
|---|---|---|---|---|---|---|---|---|
| chr4 | 36 264 381 | 36 264 382 | chr4 | 36 651 084 | 36 651 085 | i-j | inversion | |
| chr4 | 36 264 393 | 36 264 394 | chr2 | 47 652 897 | 47 652 898 | j-b | translocation | *MSH2* |
| chr2 | 61 688 253 | 61 688 254 | chr4 | 45 662 721 | 45 662 722 | b-k | translocation | *USP34* |
| chr4 | 36 651 087 | 36 651 088 | chr2 | 62 454 956 | 62 454 957 | k-d | translocation | |
| chr2 | 149 362 252 | 149 362 253 | chr4 | 55 202 616 | 55 202 617 | g-m | translocation | *MBD5* |
| chr2 | 61 748 483 | 61 748 484 | chr2 | 149 116 864 | 149 116 865 | f-c | inversion | *XPO1, MBD5* |
| chr2 | 6 168 824 | 6 168 825 | chr2 | 149 116 864 | 149 116 865 | c-g | inversion | *USP34, MBD5* |
| chr2 | 47 652 886 | 47 652 887 | chr2 | 70 215 999 | 70 216 000 | a-e | inversion | |
| chr2 | 62 454 958 | 62 454 959 | chr4 | 45 700 661 | 45 700 662 | d-l | translocation | |
| chr4 | 49 157 066 | 49 157 067 | chr2 | 149 534 002 | 149 534 003 | l-h | translocation | *EPC2, MBD5* |
