## Supplementary material for "Zooming into rearranged genome: applying pipeline of cytological, genomic, and transcriptomic methods for structural variant interpretation": Results of CNV prediction tools

**Supplementary Table 3**

### Results of CNV prediction tools

| Chromosome | State | gatk_start | gatk_end | gatk_GTinfo | cnvkit_start | cnvkit_end | conifer_start | conifer_end | consensus_start | consensus_end | consensus_range | Genes | OMIM_genes_disease_causing | Overlaps_DGV | DGV_hits |
|---|---|---|---|---|---|---|---|---|---|---|---|---|---|---|---|
| chr16 | dup | 2553131 | 2555974 | ./.:3:8:44:93:3:43 | 97433 | 17202041 | 2553464 | 2569420 | 2553464 | 2555974 | 2510 | TBC1D24 | TBC1D24(AD) | False | [] |
| chr22 | dup | 39381557 | 39388749 | ./.:3:5:15:31:20:6 | 27073039 | 51220728 | 39378404 | 39388499 | 39381557 | 39388499 | 6942 | APOBEC3B | none | True | [(39356511, 39395495, 'dup'), (39378404, 39388783, 'dup')] |
| chr12 | dup | 96394515 | 96429556 | ./.:4:19:53:85:13:54 | 96394764 | 96429306 | 96388566 | 96617559 | 96394764 | 96429306 | 34542 | LTA4H,ENSG00000257878 | none | False | [] |
| chr11 | dup | 818633 | 837054 | ./.:3:20:2:59:3:3 | 150500 | 3818563 | 830233 | 838044 | 830233 | 837054 | 6821 | CRACR2B,CD151,ENSG00000255108 | CD151(AR) | False | [] |
| chr11 | dup | 818633 | 837054 | ./.:3:20:2:59:3:3 | 150500 | 3818563 | 822378 | 829549 | 822378 | 829549 | 7171 | PNPLA2,CRACR2B,ENSG00000255108 | PNPLA2(AR) | False | [] |
| chr11 | dup | 725960 | 760526 | ./.:3:8:12:64:9:12 | 150500 | 3818563 | 726881 | 767368 | 726881 | 760526 | 33645 | EPS8L2,ENSG00000269915,ENSG00000303334,TALDO1 | EPS8L2(AR),TALDO1(AR) | False | [] |
| chr5 | dup | 180375055 | 180430038 | ./.:4:7:5:24:19:11 | 179347313 | 180582938 | 180376894 | 180420178 | 180376894 | 180420178 | 43284 | BTNL8,ENSG00000299843,BTNL3 | none | True | [(180359732, 180442908, 'dup'), (180368242, 180418332, 'dup'), (180417194, 180491462, 'dup'), (180372155, 180491462, 'dup'), (180395230, 180432024, 'dup')] |
