## Supplementary material for "Zooming into rearranged genome: applying pipeline of cytological, genomic, and transcriptomic methods for structural variant interpretation": Nanopore sequencing coverage date (FC - fold change)

**Supplementary Table 4**

### Nanopore sequencing coverage date (FC - fold change)

| Target region | WGS | | Adaptive sampling | | nCATS | |
|---|---|---|---|---|---|---|
| | Average coverage within target region | FC over average genome coverage | Average coverage within target region | FC over average genome coverage | Average coverage within target region | FC over average genome coverage |
| chr2: 47620000-47680000 | 0.30 | 0.23 | 1.03 | 1.78 | 1.50 | 2.17 |
| chr2: 61655000-61720000 | 2.32 | 1.78 | 1.06 | 1.83 | 0.73 | 1.06 |
| chr2: 62425000-62485000 | 1.36 | 1.05 | 1.43 | 2.47 | 4.28 | 6.20 |
| chr2: 70185000-70245000 | 1.60 | 1.23 | 0.85 | 1.47 | 1.13 | 1.64 |
| chr2: 149320000-149385000 | 0.54 | 0.42 | 1.02 | 1.76 | 0.156 | 0.23 |
| chr4: 36230000-36295000 | 0.95 | 0.73 | 0.67 | 1.16 | 8.25 | 11.96 |
| chr4: 45630000-45740000 | 0.96 | 0.74 | 1.14 | 1.97 | 4.90 | 7.10 |
| chr4: 55170000-55270000 | 1.19 | 0.92 | 0.22 | 0.38 | 0.34 | 0.49 |
