## Supplementary material for "Zooming into rearranged genome: applying pipeline of cytological, genomic, and transcriptomic methods for structural variant interpretation": Regions of the genome near SV breakpoints with outlier compartment scores and the corresponding gene content

**Supplementary Table 5**

### Regions of the genome near SV breakpoints with outlier compartment scores and the corresponding gene content

| Cchromosome | Start | End | Genes | OMIM_genes_disease_causing |
|---|---|---|---|---|
| chr2 | 3350000 | 3400000 | TRAPPC12, EIPR1 | |
| chr2 | 51950000 | 52000000 | NRXN1-DT | |
| chr2 | 52250000 | 52300000 | none | |
| chr2 | 61100000 | 61150000 | REL-DT, REL | |
| chr2 | 68350000 | 68400000 | DNAAF10, PNO1 | |
| chr2 | 71250000 | 71300000 | OR7E91P, NAGK | |
| chr4 | 1300000 | 1350000 | MAEA, UVSSA | UVSSA (AR) |
| chr4 | 34800000 | 34850000 | none | |
| chr4 | 36250000 | 36350000 | DTHD1 | |
| chr4 | 37650000 | 37700000 | RELL1 | |
| chr4 | 41950000 | 42000000 | TMEM33, SLC30A9 | |
| chr4 | 42650000 | 42700000 | ATP8A1 | |
| chr4 | 45650000 | 45700000 | none | |
| chr4 | 47450000 | 47500000 | COMMD8, ATP10D | |
| chr4 | 47900000 | 47950000 | NFXL1, CNGA1 | CNGA1 (AR) |
| chr4 | 48800000 | 48850000 | OCIAD1 | |
| chr4 | 57250000 | 57350000 | AASDH, PPAT, PAICS, SRP72 | |
| chr4 | 57750000 | 57850000 | REST, NOA1, POLR2B | |
| chr4 | 60050000 | 60100000 | none | |
| chr4 | 186300000 | 186350000 | LRP2BP, UFSP2, CFAP96 | UFSP2 (AR) |
