## Supplementary material for "Zooming into rearranged genome: applying pipeline of cytological, genomic, and transcriptomic methods for structural variant interpretation": List of differentially expressed genes identified in the patient-derived iPSCs

**Supplementary Table 6**

### List of differentially expressed genes identified in the patient-derived iPSCs

| Gene symbol | Gene | logFC | padj | Chromosome | Start | End |
|---|---|---|---|---|---|---|
| | | | UP REGULATED | | | |
| HAND1 | ENSG00000113196.3 | 6.48879598072741 | 2.4512327873229216 | chr5 | 153854529 | 153857916 |
| **CXCL5** | **ENSG00000163735.7** | **3.52915746325623** | **3.8502446885135497** | **chr4** | **74861359** | **74864394** |
| CST1 | ENSG00000170373.8 | 4.03950977781056 | 4.75353266783989 | chr20 | 23728199 | 23731905 |
| ZNF662 | ENSG00000182983.15 | 2.70587954752645 | 6.362198837072051 | chr3 | 42947433 | 42960826 |
| FLG-AS1 | ENSG00000237975.7 | 4.35231477855618 | 3.422057682833924 | chr1 | 152309544 | 152339168 |
| LINC00707 | ENSG00000238266.2 | 2.89670984275252 | 3.9571521715829996 | chr10 | 6821560 | 6884868 |
| PPIAP46 | ENSG00000260266.1 | 2.51840010712666 | 2.3149451960448832 | chr15 | 74653893 | 74654382 |
| AC005670.2 | ENSG00000262633.2 | 6.31427171543764 | 3.8345678763028195 | chr17 | 45000499 | 45124520 |
| PNMA6B | ENSG00000268883.2 | 9.46766558303415 | 7.288854717638465 | chrX | 152244152 | 152245351 |
| NA | ENSG00000288597.1 | 3.00996492102607 | 4.75353266783989 | chrX | 102961962 | 103173630 |
| | | | DOWN REGULATED | | | |
| SEC31B | ENSG00000075826.17 | -2.16187539509663 | 2.676736096604476 | chr10 | 102246396 | 102279621 |
| MYH14 | ENSG00000105357.19 | -3.55022165029944 | 13.803423821797358 | chr19 | 50691443 | 50813802 |
| SEC16B | ENSG00000120341.18 | -3.23581414253853 | 2.6662503864879157 | chr1 | 177893091 | 177953438 |
| EDA2R | ENSG00000131080.15 | -2.31997197247855 | 4.266075216707363 | chrX | 65814226 | 65859140 |
| KCNC3 | ENSG00000131398.15 | -3.08438932485316 | 9.2432665636929 | chr19 | 50815194 | 50836772 |
| GSTM1 | ENSG00000134184.13 | -9.82135370808125 | 6.523148989992148 | chr1 | 110229962 | 110251661 |
| **ANKRD36** | **ENSG00000135976.20** | **-2.01937395262366** | **4.871237192455953** | **chr2** | **97778890** | **97930258** |
| LGI4 | ENSG00000153902.14 | -2.386156893246 | 4.991675991482525 | chr19 | 35615414 | 35633355 |
| ADAMTS13 | ENSG00000160323.19 | -2.18542200574465 | 5.942220941854891 | chr9 | 136279478 | 136324524 |
| CCDC78 | ENSG00000162004.18 | -2.1615045599495 | 4.731649744548168 | chr16 | 772581 | 776954 |
| PABPC5 | ENSG00000174740.8 | -4.52143008881209 | 2.8981858562515255 | chrX | 90689594 | 90693583 |
| GOLGA8A | ENSG00000175265.17 | -2.17610289160929 | 3.588747113197165 | chr15 | 34671136 | 34730052 |
| MAPK15 | ENSG00000181085.15 | -2.35230193409399 | 3.4890741627540254 | chr8 | 144798449 | 144804628 |
| POU3F4 | ENSG00000196767.9 | -4.71187867441003 | 4.296038878852289 | chrX | 82763298 | 82767135 |
| CCDC180 | ENSG00000197816.16 | -2.63611113210479 | 5.284043136361207 | chr9 | 100069586 | 100141033 |
| LAT | ENSG00000213658.12 | -2.35727177731166 | 2.330816474837783 | chr16 | 28996129 | 29002105 |
| MEG3 | ENSG00000214548.18 | -12.9220072676572 | 14.231369589688013 | chr14 | 101292455 | 101327363 |
| BX322639.1 | ENSG00000215146.5 | -12.2622858023359 | 10.705712761423573 | chr10 | 42827314 | 42863422 |
| MEG9 | ENSG00000223403.6 | -7.83171864264814 | 8.043003984021174 | chr14 | 101509245 | 101539274 |
| MEG8 | ENSG00000225746.13 | -9.54820601687755 | 6.262947114440012 | chr14 | 101245543 | 101327368 |
| SAPCD1 | ENSG00000228727.9 | -2.18899163425158 | 2.6144906259286227 | chr6 | 31730433 | 31732627 |
| AL359715.1 | ENSG00000233967.7 | -2.94238679902047 | 4.75353266783989 | chr6 | 81151012 | 81172780 |
| FAM157A | ENSG00000236438.7 | -3.2390637335925 | 6.286010565645328 | chr3 | 197907626 | 197917236 |
| NPIPB5 | ENSG00000243716.10 | -2.5059386462656 | 10.705712761423573 | chr16 | 22490442 | 22547861 |
| AP006284.1 | ENSG00000254815.6 | -2.43671543938941 | 3.102642218850376 | chr11 | 557595 | 560114 |
| AL512274.1 | ENSG00000261068.2 | -2.17118668081356 | 2.1665388512476396 | chr6 | 42059971 | 42061997 |
| MYO15B | ENSG00000266714.9 | -3.03877156183312 | 7.4841463961689705 | chr17 | 73583881 | 73622929 |
| MIR222HG | ENSG00000270069.1 | -2.03527089009161 | 3.5256851842704884 | chrX | 45604638 | 45711303 |
| ZSWIM8-AS1 | ENSG00000272589.1 | -7.61947835069866 | 3.738978339181827 | chr10 | 75556272 | 75561157 |
| CU634019.1 | ENSG00000277067.4 | -11.3146797475663 | 9.208405909874752 | chr1 | 7048748 | 7116635 |
| CHMP1B2P | ENSG00000278530.4 | -6.24105352743836 | 3.2511653773726064 | chrX | 79527564 | 79590726 |
| CU633906.2 | ENSG00000278903.3 | -2.52450003498386 | 4.097680959464776 | chr21 | 6318370 | 6360415 |
| AP000866.6 | ENSG00000279342.2 | -2.2288718551279 | 2.867587367689957 | chr11 | 124659136 | 124662714 |
| AC010507.2 | ENSG00000282339.1 | -2.16707727610151 | 2.8022439660778224 | chr19 | 200499 | 201174 |
| AC092299.1 | ENSG00000282416.1 | -2.80772798756638 | 5.158674841987909 | chr19 | 197310 | 198066 |
| LINC01002 | ENSG00000282508.2 | -2.61147946210937 | 4.7118387604466765 | chr19 | 181465 | 246555 |
